# Structural Basis of Second-Generation HIV-1 Maturation Inhibitor Activity by Magic Angle Spinning NMR

**DOI:** 10.64898/2026.11.05.764638

**Authors:** Kiran Kumar, Roman Zadorozhnyi, Timothy N. Porter, Jayden A. Martinez, Caitlin M. Quinn, Sherimay D. Ablan, Eric O. Freed, Theodore J. Nitz, Carl T. Wild, Barbie K. Ganser-Pornillos, Owen Pornillos, Angela M. Gronenborn, Tatyana Polenova

## Abstract

The CA-SP1 junction of HIV-1 Gag forms a six-helix bundle (6HB) within the immature Gag lattice, sequestering the CA-SP1 cleavage site. Maturation inhibitors (MIs), such as Bevirimat (BVM) and its analogs, are an emerging class of HIV-1 antiviral compounds that stabilize Gag maturation intermediates by interfering with CA-SP1 proteolytic processing. Here, we report the binding modes of a potent second-generation BVM analog with an extended scaffold, HRF-4467, in crystalline CA_CTD_-SP1-NC_11_ assemblies with inositol hexakisphosphate (IP6), as determined by magic-angle-spinning solid-state NMR (MAS NMR) spectroscopy. The atomic-level structure reveals that HRF-4467 binds asymmetrically in the SP1-NC11 six-helix bundle and stabilizes its helical conformation in the immature lattice, in both the wild-type and the BVM-resistant SP1-V7A Gag variant. Additionally, in the presence of HRF-4467, IP6 adopts distinct conformations in the wild-type CA_CTD_-SP1-NC_11_ and the SP1-V7A variant. Taken together, our results establish a molecular basis for HRF-4467’s improved antiviral activity relative to BVM and suggest strategies for designing inhibitors with enhanced performance against MI-resistant HIV-1 variants for potential therapeutic use.

## INTRODUCTION

During HIV-1 maturation, proteolytic processing of the Gag polyprotein by the viral protease triggers extensive structural rearrangements that convert newly assembled, non-infectious particles into mature, infectious virions^1–4^. Virion assembly is driven by the 55-kDa Gag polyprotein precursor, which contains the structural domains required to form immature virus particles and initially assembles at the inner leaflet of the plasma membrane of infected cells^5^. Following budding, Gag processing occurs in a tightly orchestrated sequence of steps, generating its constituent mature Gag proteins – matrix (MA), capsid (CA), nucleocapsid (NC), and p6 – along with two short spacer peptides, SP1 and SP2^6^. These proteolytic events drive major structural rearrangements within the virion, including reorganization of the immature Gag lattice into the mature CA lattice^7, 8^. The final step of Gag processing is the cleavage of SP1 from the CA-SP1 maturation intermediate^6, 9^. Disruption or slowing of CA-SP1 cleavage interferes with viral maturation, resulting in non-infectious virions^10, 11^. Small-molecule maturation inhibitors (MIs) that specifically target the CA-SP1 cleavage site therefore represent a promising strategy for HIV-1 antiviral therapy.

Previous studies have shown that slow cleavage at the CA-SP1 junction results from structural sequestration of the proteolytic site^10, 12^. Within the immature HIV-1 Gag lattice, the CA-SP1 junction adopts an α-helical conformation, and six CA-SP1 chains organize into a six-helix bundle (6HB) that stabilizes the Gag hexamers. The cellular metabolite inositol hexakisphosphate (IP6) further stabilizes the Gag hexamer by binding between two rings formed by the CA lysine residues K158 and K227^10, 13–15^. IP6 plays a crucial role in the viral replication cycle by promoting assembly and stabilization of the immature Gag lattice and facilitating formation and stabilization of the mature CA capsid^13^. MIs, such as Bevirimat (BVM or PA-457, **Figure 1**)^16, 17^, bind inside the 6HB, thereby stabilizing the bundle and restricting protease access to the CA-SP1 site^17, 18^, preventing the helix-to-coil transition and the subsequent proteolytic processing of Gag required for viral maturation. Other BVM-derived MIs^19–24^, as well as the structurally distinct pyridone-based MI PF-46396, similarly inhibit CA-SP1 processing and impair HIV-1 maturation^14, 25^.

**Figure 1.**
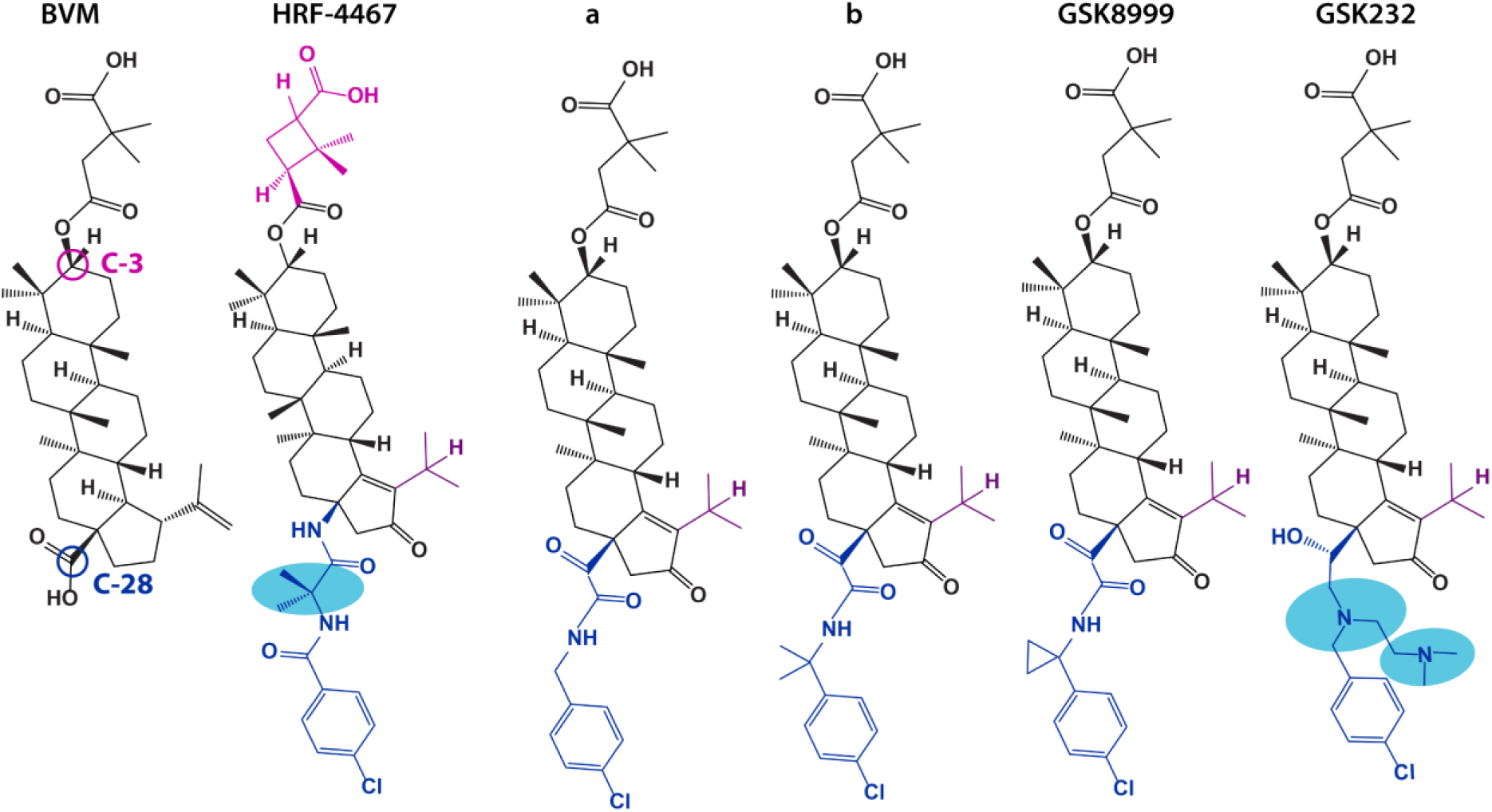
Structures of HRF-4467 and other BVM-derived HIV-1 maturation Inhibitors (MIs). The corresponding IC_50_ values of MIs are given in **Table 1**.

**Table 1.**
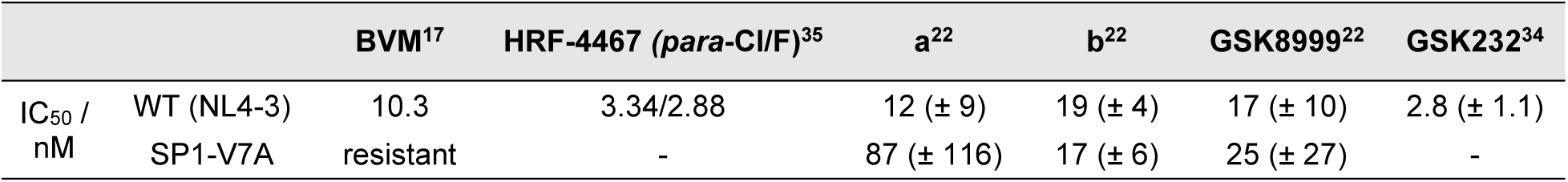
Potency of HRF-4467 and other BVM-derived maturation inhibitors against HIV-1 Gag.

The antiviral activity and molecular mechanisms of action of BVM and first-generation BVM analogs have been extensively characterized^10, 12, 26–29^. Biochemical studies have demonstrated that BVM significantly blocks CA-SP1 processing (p25 to p24) in intact virions^17^. However, naturally occurring SP1 polymorphisms in HIV-1 subtypes are associated with reduced susceptibility to BVM, particularly the SP1-V7A variant^30^. Although the parent BVM compound exhibits nanomolar antiviral activity^23, 31^, resistance frequently arises from mutations located near the CA-SP1 cleavage site. These include CA-H226Y, CA-L231F, and CA-L231M substitutions in the CA_CTD_ region; SP1-A1V, SP1-A3T, and SP1-A3V within the CA-SP1 boundary region^31, 32^; and naturally occurring polymorphisms within the SP1 QVT motif^30^.

Our previous study established the structural basis for BVM binding and resistance^10^, reporting the atomic-level structure of BVM-bound CA_CTD_-SP1 assemblies in the presence of IP6, and characterized the effects of BVM-resistant SP1-A1V and SP1-V7A Gag variants. We showed that BVM binds asymmetrically within the 6HB, inducing structural rearrangements that tighten the 6HB pore while forcing IP6 into a unique tilted conformation and arresting its dynamics^10^. We also showed that BVM stabilizes the 6HB and reduces SP1 and side-chain dynamics in the wild-type (WT) CA_CTD_-SP1 assemblies, whereas the SP1-A1V and SP1-V7A variants exhibit distinct mechanisms of BVM resistance: BVM binding is abolished or substantially weakened in SP1-A1V, whereas BVM binds to SP1-V7A but fails to induce the 6HB stabilization observed in WT assemblies^10^. These studies established the structural basis for BVM activity and resistance, and provided a structural framework for understanding MI interactions with the CA-SP1 6HB.

The emergence of BVM resistance and naturally occurring Gag polymorphisms that compromise BVM potency prompted the development of more potent second-generation MIs by modifying the BVM scaffold, particularly at the C-3 and C-28 positions, to achieve broader polymorphic coverage^19–24^. These include derivatives containing C-28 amide, alkylamine, or α-keto amide substituents and C-3 benzoic acid functionalities. For example, GSK-3532795 (BMS-955176)^19^ and its fluorinated analog GSK3640254^33^ incorporate a thiomorpholine dioxide heterocycle together with C-28 amide and C-3 benzoic acid functionalities^23^. GSK-8999 (GSK-2578999)^22^ contains a C-28 α-keto amide that incorporates a cyclopropyl group into the *para*-chlorobenzyl moiety, whereas GSK-232 (GSK2838232)^34^ contains a C-28 aminoalkyl substituent with two tertiary amines and a *para*-chlorobenzyl moiety (**Figure 1**). These modifications substantially improved antiviral potency and activity against naturally occurring Gag polymorphic variants compared with the parent BVM compound, and several second-generation MIs have advanced into clinical development^19, 23, 24, 33^. Despite their improved antiviral activity and resistance profiles (**Table 1**), the structural basis by which these modified BVM scaffolds interact with the CA-SP1 6HB and retain activity against BVM-resistant variants remains poorly understood.

The present study examines the structural basis of the second-generation MI HRF-4467 (**Figure 1 and 2a**) binding to crystalline CA_CTD_-SP1-NC_11_ assemblies. The synthesis of HRF-4467 and its biochemical and virologic characterization are described elsewhere^35^. Assemblies that include the NC_11_ region, which encompasses the first eleven residues of the NC domain, more closely recapitulate the structural context of the immature Gag lattice than assemblies that contain the shorter CA_CTD_-SP1 Gag fragment^10^. Within the immature Gag polyprotein, the NC domain is located immediately downstream of the SP1 spacer peptide^3^, placing its N-terminal residues adjacent to the SP1 tail. Structural changes at the CA-SP1 boundary during maturation may therefore influence the organization and dynamics of the adjacent SP1-NC interface^36^. Inclusion of NC_11_ also enables characterization of inhibitor interactions that extend beyond the canonical CA-SP1 6HB into the SP1-NC interface.

**Figure 2.**
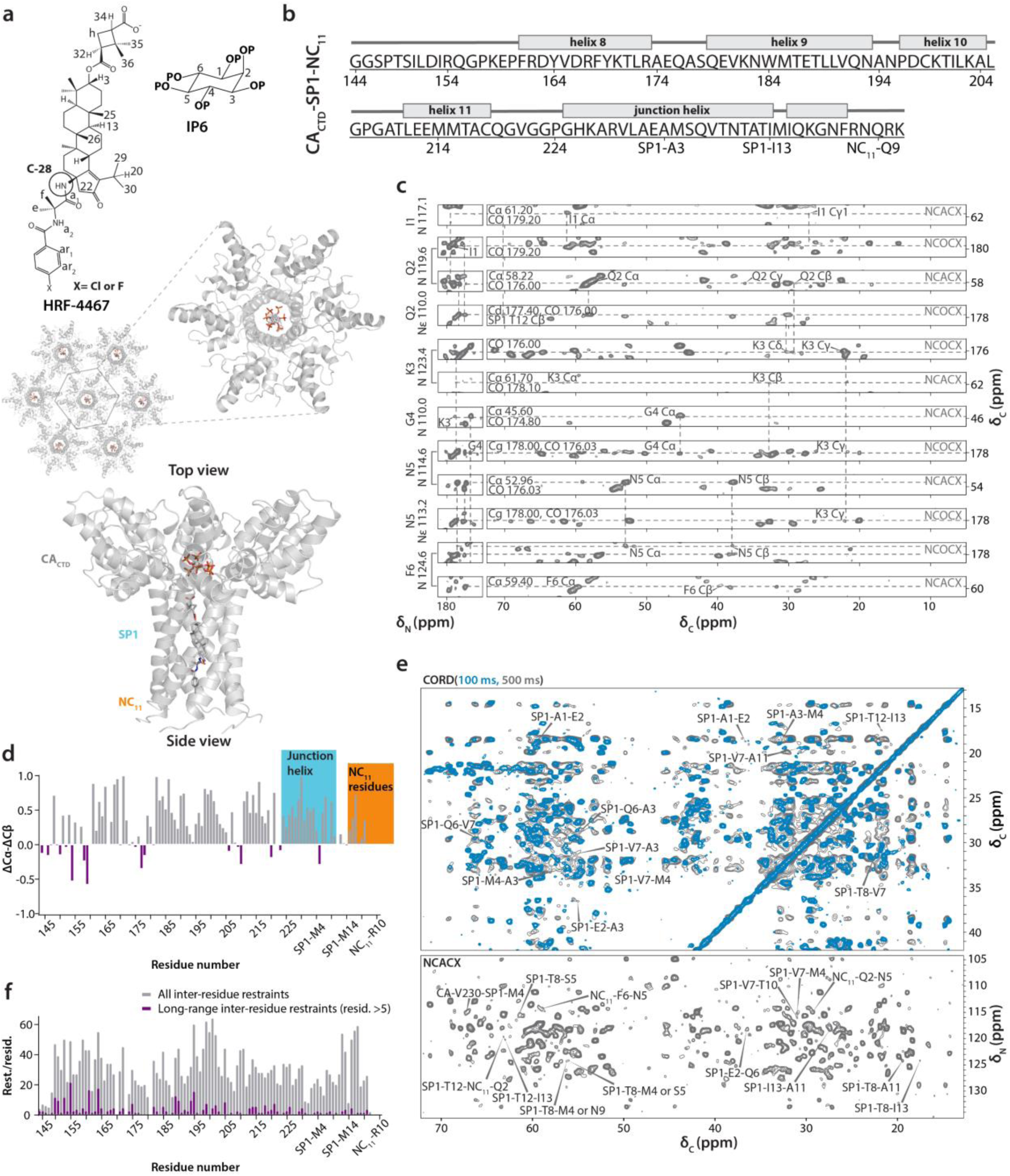
Structure of CA_CTD_-SP1-NC_11_ assemblies in the presence of HRF-4467 and IP6. **a)** Chemical structures of HRF-4467 and IP6 (top left), and the hexamer-of-hexamers (HOH) structural model: the top and side views of the central hexamer (middle and bottom left, respectively). **b)** Amino acid sequence with annotated secondary structure elements of CA_CTD_-SP1-NC_11_/HRF-4467/IP6 assemblies, based on TALOS+ analysis of the experimental chemical shifts. **c)** Representative strips of 3D NCACX and NCOCX spectra of CA_CTD_-SP1-NC_11_/HRF-4467/IP6 showing sequential assignments for a NC_11_ residue stretch I1-F6. **d)** Secondary chemical shift analysis showing the structure propensity of all residues, with junction helix and NC_11_ residues highlighted in blue and orange, respectively. **e)** Overlay of selected regions of 2D CORD spectra of CA_CTD_-SP1-NC_11_/HRF-4467/IP6 assemblies acquired with CORD mixing times of 100 ms (blue) and 500 ms (grey), together with selected regions of 2D NCACX spectrum. **f)** Number of all inter-residue and long-range MAS NMR restraints per residue plotted *vs.* residue number.

HRF-4467 shares structural features with the clinically investigated MIs GSK-8999 and GSK-232, as well as related compounds a and b shown in **Figure 1**, including a chlorophenyl moiety, while incorporating distinct isobutyramide and chlorobenzamide functionalities. Additional modifications relative to the parent BVM scaffold include cyclization of the C-3 substituent to a 2,2-dimethylcyclobutane moiety and an α,β-unsaturated ketone within the cyclopentenone ring. Intriguingly, HRF-4467 exhibits high antiviral potency (**Table 1**) despite the non-basic nature of its amide functionalities compared with the more basic amine functionalities of GSK-3532795 or GSK-232^19, 34^. Thus, the increased potency of HRF-4467 relative to the parent BVM cannot be attributed to its chemical properties alone, underscoring the need to understand the structural basis of its interactions with the immature Gag lattice.

Here, we report atomic-resolution structures of crystalline CA_CTD_-SP1-NC_11_ (**Figure 2b**) and the BVM-resistant CA_CTD_-SP1(V7A)-NC_11_ variant, bound to HRF-4467 in the presence of the assembly cofactor IP6, determined by MAS NMR spectroscopy. Our results reveal that HRF-4467 binds within the canonical CA-SP1 6HB binding pocket and extends into the SP1-NC interface, while analysis of its fluorinated analog reveals that HRF-4467(F)’s fluorophenyl moiety adopts multiple conformations upon binding. HRF-4467 binding also induces distinct IP6 conformations in WT and SP1-V7A assemblies. The atomic-resolution structural models provide a molecular basis for HRF-4467 activity against the BVM-resistant SP1-V7A variant, suggesting that the extended BVM scaffold and lipophilicity imparted by the isobutyramide moiety enhance hydrophobic interactions within the 6HB, thereby reducing dependence on the SP1-V7 side chain. Overall, our findings establish a structural framework for designing more potent next-generation MIs.

## EXPERIMENTAL

### Protein expression and purification

Uniformly ^13^C/^15^N-labeled CA_CTD_-SP1-NC_11_ was expressed in E. *coli* grown in M9 minimal medium containing ^13^C-glucose and ^15^N-ammonium chloride as the sole carbon and nitrogen sources. A 50-mL seed culture was prepared in M9 medium supplemented with 5× M9 salts, trace elements, MgSO_4_, CaCl_2_, thiamin, biotin, and kanamycin, inoculated from a single colony, and grown overnight at 37 °C with shaking at approximately 250 rpm. The seed culture was used to inoculate 2 L of identically supplemented M9 medium containing 4 g ^13^C-glucose and 2 g ^15^N-ammonium chloride. Cells were grown at 37 °C with shaking to an OD600 of 1.0–1.2, protein expression was induced with 1 mM IPTG, and cultures were incubated overnight at 20 °C. Cells were harvested by centrifugation at 5,000 × g for 10 min and either processed immediately or stored at −80 °C.

For purification, the cell pellet from a 2-L culture was resuspended in 50 mL of 50 mM Tris, pH 8.3, 1 M LiCl supplemented with two Roche protease inhibitor tablets and lysozyme, and incubated on ice for 20 min. 300 µL of a 10% solution of deoxycholate was then added, and the cells were lysed by two 20-s rounds of sonication. The lysate was clarified by centrifugation at 41,000 × g for 45 min and the supernatant was passed through a 0.45-µm filter. The clarified lysate was applied to a 5-mL HisTrap column equilibrated with 50 mM Tris, pH 8.3, 1 M LiCl, and 15 mM imidazole. After washing with five column volumes of equilibration buffer, bound protein was eluted using a linear 15–300 mM imidazole gradient over 200 mL (40 column volumes). Fractions containing CA_CTD_-SP1-NC_11_ were pooled and dialyzed overnight at 4 °C against 20 mM Tris, pH 8.0, 500 mM NaCl, and 2 mM β-mercaptoethanol. The dialyzed sample was subsequently applied to two tandem 5-mL Q anion-exchange columns equilibrated in the same buffer. CA_CTD_-SP1-NC_11_ was collected in the flow-through and column wash fractions, pooled, and concentrated to approximately 10 mg/mL. All chromatography steps were performed at 4 °C.

### Preparation of inhibitor-protein assemblies for solid-state NMR

CA_CTD_-SP1-NC_11_ assemblies were prepared by combining purified protein with inositol hexakisphosphate (IP6) and MI at equimolar concentrations. Protein concentrations typically ranged from approximately 750 µM to 1 mM, and IP6 and HRF-4467 concentrations were adjusted to match the molar concentration of CA_CTD_-SP1-NC_11_ in each reaction. IP6 and HRF-4467 were added from 10 mM stock solutions. Samples were mixed gently and incubated overnight at 20 °C to promote assembly. Assembly was evaluated by transmission electron microscopy. The resulting assemblies were centrifuged at 10,000 × *g* and packed into 3.2-mm and 1.9-mm Bruker rotors. For 1.3-mm Bruker rotors, the assemblies were packed at 68,300 × *g* using an ultracentrifuge rotor-packing tool.

### MAS NMR spectroscopy

All MAS NMR experiments were performed on 500 MHz Bruker Avance III, 600 MHz Bruker Avance III HD, and 850 MHz Bruker Avance III spectrometers. 2D CORD, NCOCX, and NCACX experiments used for chemical-shift assignments were acquired on the 850 MHz spectrometer using a 3.2 mm E-free HCN probe at the MAS frequency of 14 kHz, controlled to within ± 10 Hz by a Bruker MAS-3 controller. The temperature for all experiments was calibrated using the ^79^Br T_1_ relaxation time in KBr powder^37^.

The 2D rINEPT-TOBSY and 3D NCOCX and NCACX experiments were acquired on the 600 MHz spectrometer using a 3.2 mm E-free HCN probe at the MAS frequency of 14 kHz, controlled to within ± 10 Hz by a Bruker MAS-3 controller. Typical π/2 pulses were 2.5-2.6 μs for ^1^H, 3.0-3.6 μs for ^13^C, and 4.6-4.7 μs for ^15^N. For ^15^N/^13^C-^1^H cross-polarization (CP), contact times were 1-1.2 ms, with typical radio frequency (rf) field of 50 kHz (^15^N), 49 kHz (^13^C), and 66 kHz (^1^H), respectively. A linear 20% amplitude ramp was applied on ^1^H, with the center of the ramp matched to the n=1 Hartmann-Hahn condition. The contact time for the band-selective ^15^N-^13^C magnetization transfer was 6.5 ms, and the rf fields were 35/34 kHz (^13^C) and 21/49 kHz (^15^N) for NCA and NCO transfers, respectively. 2D R2_n_^v^ driven (CORD)^38^ spectra were obtained with 50, 100, 250, and 500 ms mixing times, and 2D NCACX spectrum was acquired with 50 ms mixing time. C9^1^_3_ J-coupling mediated 2D rINEPT-TOBSY^39, 40^ spectra were obtained with the mixing time of 7.6 ms. Spinal-64^41^ high power decoupling (80 kHz) was used during the evolution and acquisition periods. The recycle delay was 2 s. The sample temperature was set to 278 K, controlled to within ± 1 K using the Bruker temperature controller unit.

^19^F-detected 2D HF HETCOR MAS NMR spectra were acquired on a Bruker AVIII 500 MHz spectrometer outfitted with a 1.3 mm HFX probe. The typical π/2 pulse lengths were 2.3 μs (^1^H) and 2.2 μs (^19^F). The ^1^H-^19^F cross-polarization employed a linear amplitude ramp of 90-110% on ^1^H, with the center of the ramp matched to the n=1 Hartmann-Hahn condition; the typical rf fields were 35 kHz (^19^F) and 95 kHz (^1^H); the CP contact time was 1.5 ms. SWf-TPPM^42^ ^1^H decoupling (15 kHz) was applied during the acquisition. The recycle delay was 5 s and 3 s for 1D and 2D experiments, respectively. The MAS frequency was 60 kHz, controlled to within ± 10 Hz; the sample temperature was set to 303 K, controlled to within ± 5 K.

^13^C/^31^P-detected 2D HC and HP HETCOR MAS NMR spectra were acquired on a Bruker Avance III HD 850 MHz spectrometer outfitted with a 1.9 mm HCN and HX probe. The typical π/2 pulse lengths were 2.1-2.5 μs (^1^H), 3.2-3.6 μs (^13^C), and 3.5 μs (^31^P). The ^1^H-^31^P/^1^H-^13^C cross-polarization employed a linear amplitude ramp of 90-110% on ^1^H, with the center of the ramp matched to the n=1 Hartmann-Hahn condition; the CP contact time was 3.5 ms for ^1^H-^31^P transfer, and 10.0 ms for ^1^H-^13^C transfer. SWf-TPPM ^1^H decoupling (12.5-15 kHz) was applied during the acquisition. The recycle delay was 2 and 1.5 s, respectively. The MAS frequency was 40 kHz, controlled to within ± 10 Hz; the sample temperature was set to 278 K, controlled to within ± 1 K. ^1^H-detected 2D CP-(H)NH and rINEPT-(H)NH MAS NMR spectra were acquired on Bruker Avance III 850 and Bruker Avance III HD 600 MHz spectrometers outfitted with a 1.3 mm HCN probe. Typical π/2 pulses were 2.5 μs (^1^H), 3.6 μs (^13^C), and 3-4.5 μs (^15^N). The ^1^H-^15^N/^15^N-^1^H cross-polarization employed a linear amplitude ramp of 110-90% on ^1^H, with the center of the ramp matched to the n=1 Hartmann-Hahn condition. For the forward ^1^H-^15^N and the reverse ^15^N-^1^H CP, the contact times were 1.2 ms and 0.4 ms, respectively, with typical rf fields of 35-40 kHz (^15^N) and 100-105 kHz (^1^H). The recycle delay was 1.2 s for both experiments. The MAS frequency was 60 kHz, controlled to within ± 10 Hz; the sample temperature was set to 303 K controlled to within ± 5 K.

In all spectra, the ^15^N and ^13^C chemical shifts were referenced to the external standards of ammonium chloride, adamantane, respectively. ^19^F chemical shifts were referenced to the high-field 2-CF_3_ resonance of mefloquine at 8.8 ppm^43^. ^1^H chemical shifts were indirectly referenced to the ^13^C chemical shifts of adamantane. ^31^P was referenced with respect to the phosphorous resonance of 85% H_3_PO_4_ at 0 ppm.

All MAS NMR spectra were processed using Bruker TopSpin and NMRPipe^44^ and analyzed in CcpNmr analysis software^45^. Each 2D and 3D spectrum was processed by applying 30°, 60°, and 90° shifted sine bell apodization, followed by a Lorentzian-to-Gaussian transformation in all dimensions. Two-fold forward linear prediction was used in the indirect dimension in some data sets, with zero filling applied, followed by Fourier transformation.

### Computational analysis and force field parameterization of HRF-4467

All quantum mechanical (QM) calculations were performed in Gaussian 16^46^. The starting structures of HRF-4467 (*para-*Cl and *para-*F analogs) were geometry optimized at the MP2/6-31G* level of theory using an implicit water solvation model. These geometry-optimized structures were used for force field parameterization with the CHARMM General Force Field (CGenFF) server^47^. Topology and parameter files were generated, including atomic partial charges and bond, angle, dihedral, and non-bonded parameters. The ^19^F isotropic magnetic shielding constant Density Functional Theory (DFT) calculations were performed on a geometry-optimized structure of the fluorinated analog, HRF-4467(F), using the gauge-including atomic orbital (GIAO) method at the B3LYP/cc-pVDZ level of theory. Relative conformational energies were determined by scanning the CNCC dihedral angle at the isobutyramide moiety (1, 2, 3, 4 atoms, **Figure S1 and Table S1**), corresponding to rotation about the CN bond, connecting the isobutyramide moiety to the five-membered ring. Conformations within 5 kcal/mol of the global minimum were selected for subsequent electrostatically embedded QM calculations. The computed isotropic magnetic shielding constants were converted to ^19^F chemical shifts using a calibration derived from 707 aromatic and aliphatic C–F groups^48^ and referenced to mefloquine on a scale where trifluoroacetic acid is set to 0 ppm, corresponding to an offset of approximately 76.5 ppm relative to CFCl_3_, and experimental ¹⁹F chemical shifts were subsequently converted to isotropic magnetic shielding constants using the calibration equation in **Table S2**. Calculated absolute magnetic shielding constants (σᵢ) were converted to chemical shifts using δᵢ = σ_ref_ − σᵢ, with σ_ref_ determined by linear regression against experimental chemical shifts. This referencing approach has also been used in several previous studies^49, 50^. To support the calibration of the calculated ^19^F chemical shifts, we collected solution ^19^F NMR spectra of free HRF-4467(F) in three solvent systems (**Figure S2**, Supporting Information).

For each of the seven selected conformations, the QM region included HRF-4467(F) and six residues spanning the SP1-A11 to NC_11_-Q2 region of the CA_CTD_-SP1-NC_11_ 6HB. The N- and C-terminal residues of each peptide fragment were capped with ACE (acetyl) and NME (N-methylamide) groups, respectively, resulting in a 772-atom QM model. The remaining residues of the 6HB and the bound IP6 molecule were represented by fixed atomic point charges, providing electrostatic embedding based on the lowest-energy MAS NMR-derived structure. For each of the seven embedded QM models, we first achieved self-consistent field (SCF) convergence using the quadratic convergence (QC) algorithm. The converged electronic wavefunctions were subsequently used to calculate ^19^F isotropic magnetic shielding constants as described above. The IP6 force field parameters were derived in our previous study^10^ and used here for structure calculations and predicted cLogP values for HRF-4467, HRF-4467(F), and GSK-232 were obtained from their chemical structures using ChemBioDraw Ultra 14.0.

### Structure calculations and refinement

The distance restraints were derived from multiple MAS NMR spectra. Both unambiguous and ambiguous restraints (up to 4-fold ambiguity) were considered. ^13^C-^13^C restraints were obtained from the 2D CORD spectra with mixing times of 50 ms, 100 ms, 250 ms, and 500 ms, and ^15^N-^13^C restraints were obtained from 2D NCACX spectra. ^13^C-^13^C/^15^N-^13^C intra- and inter-residue restraints are defined by upper-bound distance cutoffs of 6.5 Å and 7.2 Å, respectively. ^15^N/^13^C/^31^P/^19^F-^1^H restraints were obtained from 2D HETCOR spectra. ^15^N-^1^H and ^13^C-^1^H, ^31^P-^1^H, and ^19^F-^1^H restraints are defined by upper bounds of 7.5 Å, 9.2 Å, and 11.5 Å, respectively. Protein-to-IP_6_ and protein-to-HRF-4467 restraints were unambiguously assigned using 2D HETCOR spectra. Backbone angles phi (*φ*) and psi (*ψ*) were predicted from TALOS+^51^ using the experimental ^15^N and ^13^C chemical shifts.

All calculations were performed using Xplor-NIH (version 2.53)^52–54^. We first calculated an ensemble of single-chain CA_CTD_-SP1-NC_11_ structures using unambiguous experimental distance restraints. We generated the initial template of seven-hexamer (42-chain) assemblies by docking the single-chain structure into the density map, as described in our previous report, using an in-house UCSF Chimera script^10^. We calculated the final structures of HRF-4467 and IP_6_-bound assemblies using a hexamer-of-hexamers (HOH) template with protein-to-IP_6_ and protein-to-HRF-4467 distance restraints, along with the coordinates, topology, and force-field parameters of IP_6_ and HRF-4467. In addition, intra-chain restraints and hexamer-hexamer distance restraints were provided in the structure calculations.

A total of 120 structures were refined using a protocol adopted from our previous study^14^, using two identical simulated annealing runs. The high-temperature dynamics was performed for 10 ps with a time step of 1 fs, starting at 3,000 K. The initial velocities were randomized to achieve a Maxwell distribution at a starting temperature of 3000 K. The temperature was then reduced to 25 K in 25 K steps. At each temperature, we performed a 400-fs dynamics run with a 1 fs time step. During the annealing process, the hydrogen bond database^55^ and torsion angle database^56^ potential terms were used. We set standard bond angles, bond lengths, and improper terms, and used a cross-correlation probability distribution potential to enforce the grid size (boundary) of the hexamer assemblies using the reported density map^10^. The potential was restricted to N, C, CA, and O backbone atoms to ensure the density boundary did not influence side-chain conformations. After the simulated annealing runs, we performed a final energy minimization of the angles and Cartesian coordinates.

We selected the final 10 structures from the 120 calculated structures based on the lowest energy of the experimental restraining terms. We evaluated these 10 structures in the final ensemble using pairwise root mean square deviation (RMSD). **Tables S3 and S4** (Supporting Information) show summary statistics for the structures and restraints, as well as the precision and accuracy of the RMSD values, and **Figure S3** (Supporting Information) illustrates the final structural ensemble.

## RESULTS

### MAS NMR structure of HRF-4467 bound to CA_CTD_-SP1-NC_11_ assemblies

**Figure 2a** shows the 3D structure of CA_CTD_-SP1-NC_11_ in microcrystalline assemblies with HRF-4467 and IP6, illustrating a hexameric building block and the HOH motif comprising the Gag lattice, together with the binding poses of HRF-4467 and IP6. The CA_CTD_-SP1-NC_11_ structure with the bound HRF-4467 and IP6 was determined from multiple sets of two- and three-dimensional MAS NMR experiments. The corresponding spectra show high resolution and sensitivity, as illustrated in **Figure 2c**, with representative strips from the 3D NCACX and NCOCX experiments used to assign the NC_11_ resonances. The high peak intensities for the NC_11_ I1-F6 stretch of residues indicate that they are ordered. In contrast to the BVM-bound CA_CTD_-SP1 microcrystalline assemblies^10^, the junction helix extends through residue SP1-I13, while the ^13^C and ^15^N chemical shifts indicate that NC_11_ residues I1-F6 adopt a well-defined helical conformation (**Figure 2d**). Signals corresponding to the NC_11_ tail residues R7-K11 are obscured by spectral overlap and were therefore excluded from the structure calculations. Representative 2D ^13^C-^13^C CORD and NCACX spectra highlight medium- and long-range inter-residue correlations within the SP1 and NC_11_ regions (**Figure 2e**). These spectra provide many intra- and inter-residue restraints, which are used for resonance assignments and structural analysis of the SP1-NC junction. Overall, 2889 unambiguous ^13^C-^13^C, ^15^N-^13^C, ^13^C-^1^H, ^31^P-^1^H, and ^15^N-^1^H distance restraints were obtained, including 621 sequential (|i-j|=1), 798 medium-range (1<|i-j|< 4), and 197 long-range (|i-j|≥ 5), of which 26 are long-range inter-chain and 14 are long-range inter-hexamer restraints, as summarized in **Table S4** (Supporting Information). As shown in **Figure 2f**, 28 unambiguous restraints per residue on average were obtained and used in the structure calculations, yielding C-C restraint completeness of 38% at a 7.2 Å distance cut-off and 63% at a 6 Å cut-off.

As shown in the experimentally derived structure (**Figure 3a**), HRF-4467 occupies the central cavity of the CA-SP1 6HB in a well-defined binding pose within a tightly packed helical bundle. The overall hexamer architecture is preserved with six-fold symmetry, whereas the side chains lining the inhibitor-binding cavity adopt nonequivalent orientations across the six chains of hexamer, generating local asymmetry around HRF-4467. The binding poses of HRF-4467 were determined from 2D ^1^H-^19^F and ^1^H-^13^C HETCOR experiments (**Figure 3b and 3c)**. Based on the corresponding spectra, we detected 16 protein-to-HRF-4467(*para-*Cl/F) correlations, which enabled unambiguous positioning of the inhibitor within the 6HB of the CA_CTD_-SP1-NC_11_ assemblies: the *para*-chlorobenzamide ring is oriented toward the SP1 region, whereas the dimethylcyclobutane-linked carboxylate moiety faces the CA_CTD_. This orientation is supported by a unique intermolecular correlation between the methylene groups of the dimethylcyclobutane moiety and residue L231. In addition, the H3 proton of the triterpene scaffold correlates directly with SP1-A1, placing this region of the inhibitor close to the CA-SP1 junction. The triterpene core further engages in residue-specific contacts with SP1-M4, SP1-V7, and SP1-A11, whereas the amide group of HRF-4467’s chlorobenzamide moiety shows a direct intermolecular correlation with the first residue of the NC domain (NC_11_-I1), demonstrating that the inhibitor-binding interface extends beyond the canonical CA-SP1 6HB and into the SP1-NC_11_ junction.

**Figure 3.**
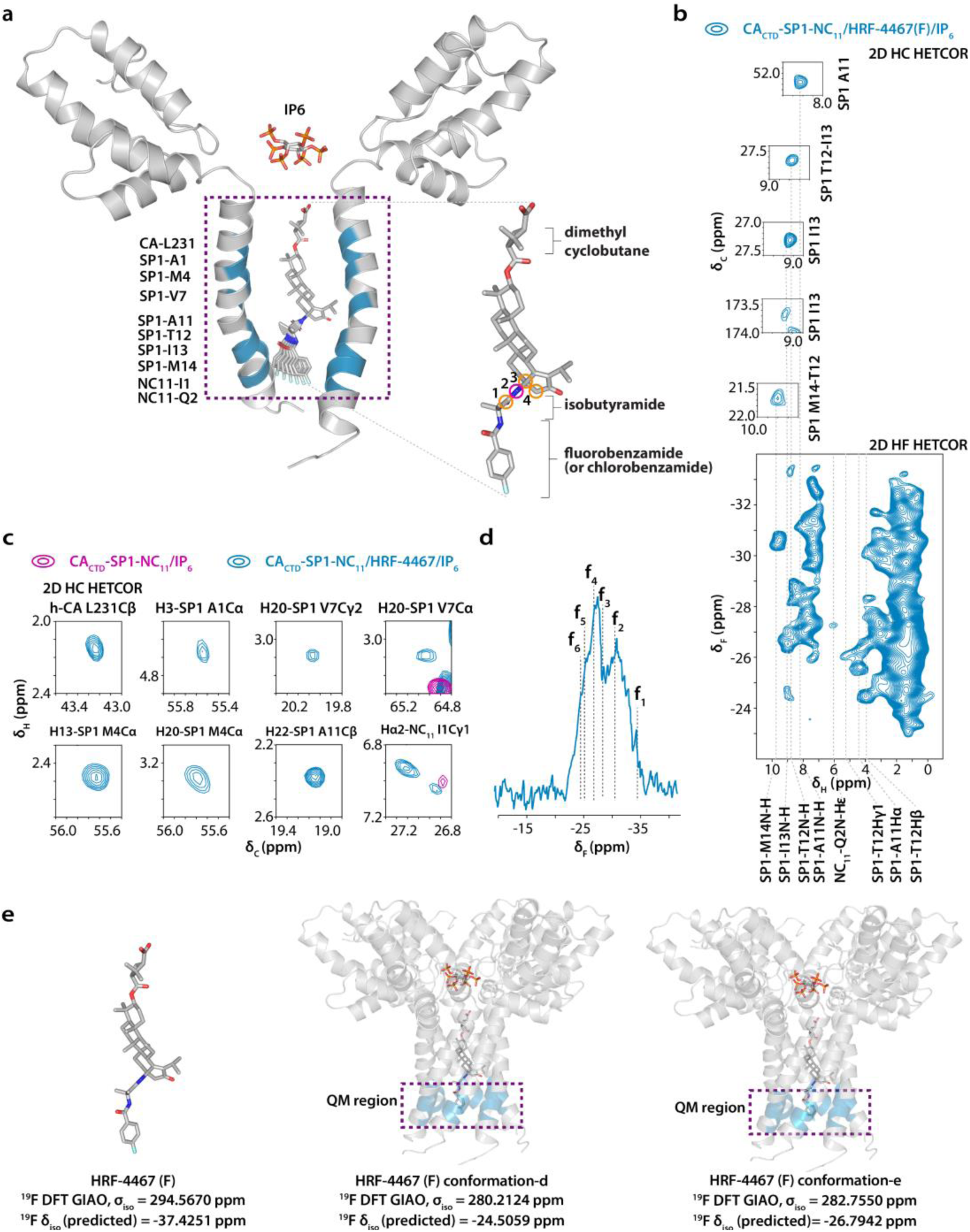
Binding poses of HRF-4467 in CA_CTD_-SP1-NC_11_/IP6 assemblies. **a)** Binding poses of HRF-4467 and IP6 in the 6HB of HIV-1 CA_CTD_-SP1-NC_11_/IP6 assemblies (two CA_CTD_-SP1-NC_11_ chains are shown, and protein-inhibitor contacts visible in NMR are highlighted). **b)** ^19^F-detected 2D HF HETCOR spectrum of CA_CTD_-SP1-NC_11_/HRF-4467(F)/IP6 assemblies. Representative expansions of ^13^C-detected HC HETCOR traces are shown as guides for assigning the corresponding HF HETCOR correlations. **c)** Selected regions of the ^13^C-detected 2D HC HETCOR spectra of CA_CTD_-SP1-NC_11_/IP6 assemblies in the absence and presence of HRF-4467, highlighting the intermolecular protein–HRF-4467 correlations. **d)** ^1^H-^19^F CPMAS spectrum of CA_CTD_-SP1-NC_11_/HRF-4467(F)/IP6 assemblies, revealing conformational heterogeneity of bound HRF-4467 within the 6HB, with (f1–f6) resonances corresponding to distinct bound conformations. **e)** Free HRF-4467(F) and two representative protein-embedded QM models selected from conformations within 5 kcal/mol of the global minimum, with their corresponding ^19^F isotropic shielding constants and calculated chemical shifts.

The experimental structure further reveals that, owing to its extended scaffold, HRF-4467 traverses the central channel of the CA-SP1 6HB and forms extensive interactions with the surrounding binding pocket. The CA-SP1 cleavage site remains buried within the tightly packed helical bundle, consistent with restricted accessibility to the viral protease. Collectively, these structural features suggest that HRF-4467 stabilizes the 6HB through extensive hydrophobic packing mediated by its triterpene core and isobutyramide moiety, together with local side-chain reorientation within the 6HB. In addition, the dimethylcyclobutane-linked carboxylate moiety is positioned near the positively charged ε-amino groups of the CA-K227 side chains, suggesting a potential electrostatic interaction with the K227 ring that may modulate IP6-K227 interactions and the conformational dynamics of IP6 within the 6HB pore. The electrostatic character of HRF-4467, particularly the negative electrostatic potential associated with the carboxylate moiety, is illustrated by the DFT-calculated electrostatic potential mapped onto an electron-density isosurface (**Figure S4**, Supporting Information). At the opposite end of the inhibitor, the *para*-fluoro or *para*-chlorobenzamide moiety extends toward the SP1-NC_11_ junction. 2D ^1^H–^19^F HETCOR spectra (**Figure 3b**) show unambiguous intermolecular correlations between HRF-4467(F) and SP1-A11, SP1-T12, SP1-I13, and NC_11_-Q2, establishing the proximity of the fluorinated aromatic moiety to these residues. Together, these interactions define an extended binding interface spanning the CA-SP1 6HB and the SP1-NC_11_ junction and provide a structural framework for understanding the antiviral activity of HRF-4467.

### Multiple conformations of HRF-4467 and IP6 orientations in CA_CTD_-SP1-NC_11_ assemblies

To further investigate the conformational preferences of the bound HRF-4467, ^1^H-^19^F cross-polarization (CPMAS) spectra were acquired for CA_CTD_-SP1-NC_11_/HRF-4467(F)/IP6 assemblies. The ^19^F signal is strong, indicating a rigid environment and, surprisingly, is very broad, spanning a chemical shift range of approximately 14.0 ppm, with two distinct intensity maxima centered at −26.8 and −30.5 ppm (**Figure 3d**). This broad chemical shift distribution indicates that the fluorinated aromatic moiety of HRF-4467 experiences multiple local environments when bound to the CA_CTD_-SP1-NC_11_ assemblies, consistent with multiple distinct conformations that are static on the timescale of the MAS NMR experiments. The experimental ^19^F chemical shifts of HRF-4467(F) bound to CA_CTD_-SP1-NC_11_/IP6 assemblies were converted to isotropic magnetic shielding constants using a linear calibration derived from the ^19^F chemical shifts of 707 aromatic and aliphatic C–F groups (**Table S2**)^48^, allowing for comparison with the QM models described below.

To understand the conformational distribution of bound HRF-4467, we computed the relative energies and ^19^F isotropic magnetic shielding constants of different conformers of HRF-4467(F) generated by rotating the isobutyramide moiety around the CCNC dihedral angle defined by atoms 1-2-3-4 (**Figure 3a** and **Figure S1**, Supporting Information). **Figure S1** shows the conformational landscape, with the relative energy difference (ΔE) plotted as a function of the CCNC dihedral angle. Conformations within 5 kcal/mol of the global minimum, including those at the boundaries, were selected for subsequent ^19^F isotropic magnetic shielding constant calculations, with and without the protein QM region (**Table S1**, Supporting Information). The protein-embedded calculations yielded predicted ^19^F chemical shifts of approximately −24 to −28 ppm (**Figure 3e and Table S1**, Supporting Information), consistent with the higher-frequency region of the experimentally observed ^19^F chemical shift distribution (see below) and corresponding to one of the conformations adopted by bound HRF-4467(F). The lower-frequency signals may reflect additional bound conformations or local environments not captured by the calculated models; we cannot yet assign them to specific conformations.

We next assessed the bound conformation and dynamics of IP6 in CA_CTD_-SP1-NC_11_/HRF-4467/IP6 assemblies. Comparison of the ^31^P direct-polarization (DPMAS) and CPMAS spectra (**Figure 4a**) revealed a pronounced change in IP6 dynamics upon HRF-4467 binding. In the absence of HRF-4467, IP6 resonances were clearly observed in the DPMAS spectrum but absent in the CPMAS spectrum, indicating that IP6 undergoes substantial molecular motion that averages the ^1^H-^31^P dipolar interaction and hampers cross-polarization transfer. In contrast, upon HRF-4467 binding, strong ^31^P CPMAS signals emerge, consistent with reduced IP6 dynamics on the microsecond timescale and increased ordering of the cofactor within the CA-SP1 6HB. The observed correlations between the protein and IP6 in 2D ^1^H-^31^P HETCOR spectra (**Figure 4b)** place IP6 in close proximity to CA_CTD_ residues E159, K158, P157, G223, P224, and K227, consistent with the previously observed IP6 location within the central pore of the CA-SP1 assemblies^10, 13, 29^. Inside the 6HB pore, IP6 adopts a distinct asymmetrical geometry, defined by 26 protein-to-IP6 intermolecular correlations that establish its position and conformation within the assembly. Compared with the HRF-4467-bound assembly, IP6 occupies a higher position in the pore and is substantially more tilted in the BVM-bound WT structure^10^. These observations suggest that HRF-4467 binding preserves the overall IP6-binding cavity while altering specific protein-IP6 interactions within the pore. However, the present data do not unambiguously distinguish whether IP6 adopts a single or multiple conformations. Nevertheless, the emergence of ^31^P CP signals upon HRF-4467 binding indicates that the inhibitor restricts IP6 motion and increases its ordering within the 6HB pore.

**Figure 4.**
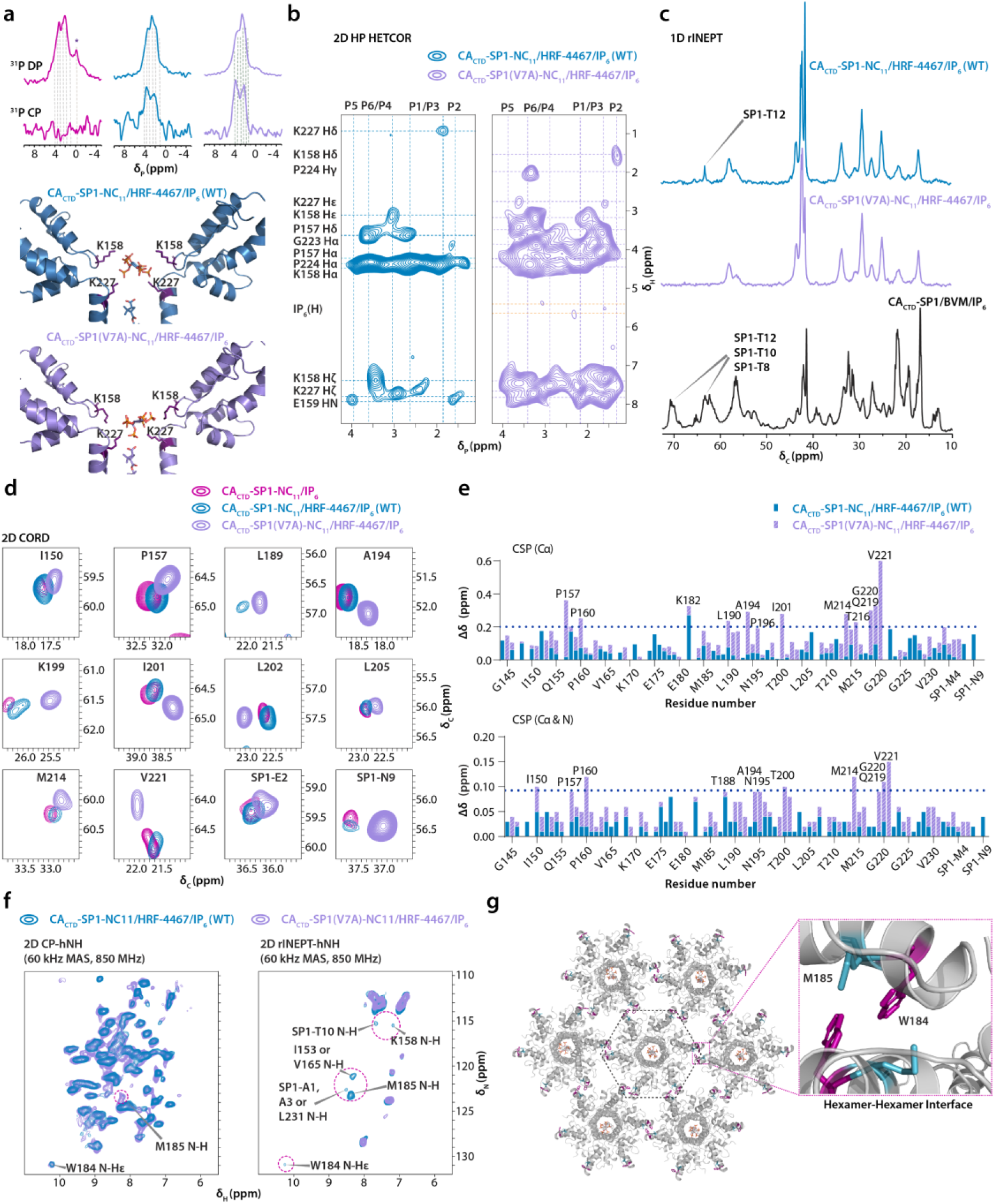
Effect of HRF-4467 binding and distinct orientations of IP6 in wild type (WT) and SP1-V7A variant of CA_CTD_-SP1-NC_11_ assemblies. **a)** ^31^P direct-polarization (DPMAS) and cross-polarization (CPMAS) spectra of CA_CTD_-SP1-NC_11_/IP6 (magenta), CA_CTD_-SP1-NC_11_/HRF-4467/IP6 (blue), and CA_CTD_-SP1(V7A)-NC_11_/HRF-4467/IP6 (violet) assemblies (top left) and distinct orientations of IP6 in the CA_CTD_-SP1-NC_11_/HRF-4467/IP6 and CA_CTD_-SP1(V7A)-NC_11_/HRF-4467/IP6 assemblies (bottom left). **b)** 2D ^1^H-^31^P correlation NMR spectra of IP6-bound CA_CTD_-SP1-NC_11_/HRF-4467/IP6 (blue), and CA_CTD_-SP1(V7A)-NC_11_/HRF-4467/IP6 (violet) assemblies. **c)** 1D rINEPT spectra of CA_CTD_-SP1-NC_11_/HRF-4467/IP6 (blue), CA_CTD_-SP1(V7A)-NC_11_/HRF-4467/IP6 (violet), and CA_CTD_-SP1/BVM/IP6 (black) assemblies. **d)** Overlay of selected regions of 2D CORD spectra of CA_CTD_-SP1-NC_11_/IP6 (magenta), CA_CTD_-SP1-NC_11_/HRF-4467/IP6 (blue), and CA_CTD_-SP1(V7A)-NC_11_/HRF-4467/IP6 (violet) assemblies, **e)** Chemical shift perturbations induced by HRF-4467 binding in CA_CTD_-SP1-NC_11_/HRF-4467/IP6 and CA_CTD_-SP1(V7A)-NC_11_/HRF-4467/IP6 assemblies plotted against residue number. **f), g)** Selected region of 2D CP and rINEPT-(H)NH spectra of CA_CTD_-SP1-NC_11_/HRF-4467/IP6 (blue) and CA_CTD_-SP1(V7A)-NC_11_/HRF-4467/IP6 (violet) assemblies showing the effect of HRF-4467 binding on WT & SP1-V7A variant spectra and structural representation of hexamer-hexamer interface region with residues M185 (purple) and W184 (blue) highlighted.

### Conformation and orientations of HRF-4467 and IP6 in CA_CTD_-SP1(V7A)-NC_11_ BVM-resistant variant assemblies

HRF-4467 binding and the associated changes in IP6 conformation are accompanied by reduced protein dynamics, as indicated by enhanced dipolar-transfer efficiency in 2D ^13^C-^13^C CORD spectra of the HRF-4467-bound WT and SP1-V7A CA_CTD_-SP1-NC_11_/IP6 assemblies compared with BVM-bound CA_CTD_-SP1/IP6 and inhibitor-free CA_CTD_-SP1-NC_11_/IP6 assemblies (**Figure S5**, Supporting Information). These spectra also indicate that HRF-4467-bound SP1-V7A assemblies are globally more rigid than the corresponding WT assemblies. To further examine the effects of HRF-4467 on protein dynamics, we acquired one-dimensional ^13^C rINEPT spectra, which selectively detect highly dynamic residues^39, 40^, for the HRF-4467-bound WT and SP1-V7A CA_CTD_-SP1-NC_11_/IP6 assemblies and the BVM-bound CA_CTD_-SP1/IP6 assemblies. Differences in rINEPT signal intensities reveal distinct dynamic behavior in the SP1-tail region (**Figure 4c**). Two-dimensional ^13^C-^13^C rINEPT-TOBSY spectra further show efficient scalar-based magnetization transfer within the SP1-tail residues in the BVM-bound assemblies (**Figure S6a**, Supporting Information), whereas the corresponding signals are strongly attenuated in the HRF-4467-bound WT and SP1-V7A assemblies (**Figure S6b**, Supporting Information). The residual SP1-T12 signal remains detectable in the HRF-4467-bound WT assemblies but is further attenuated in the SP1-V7A assemblies, indicating reduced dynamics of the C-terminal region of the junction helix in the variant. In addition to these dynamic changes, overlays of 2D ^13^C-^13^C CORD spectra reveal pronounced residue-specific chemical-shift perturbations relative to the inhibitor-free WT CA_CTD_-SP1-NC_11_/IP6 assemblies (**Figure 4d**). Quantitative analysis shows that these perturbations extend beyond the SP1 region and affect residues near the IP6-binding site (P157, A194, G220, and V221), within the 6HB (SP1-E2 and SP1-N9), and in helices 9–11 (**Figure 4e**). The perturbation patterns exhibit both shared and distinct features in the HRF-4467-bound WT and SP1-V7A assemblies, indicating similarities and differences in local environments extending from the inhibitor-binding pocket to the IP6-binding region and surrounding CA_CTD_ helices. Together, these results indicate that HRF-4467 quenches SP1-tail dynamics and stabilizes the 6HB, with more pronounced rigidification in the BVM-resistant SP1-V7A variant.

^1^H-detected ^1^H-^15^N HETCOR spectra reveal pronounced perturbations in the Q155 side-chain resonances of the HRF-4467-bound WT and SP1-V7A CA_CTD_-SP1-NC_11_/IP6 assemblies (**Figure S6c**, Supporting Information). In both assemblies, the Q155 side-chain ^15^N resonances shift from approximately 109 to 107 ppm relative to the BVM-bound CA_CTD_-SP1/IP6 and inhibitor-free CA_CTD_-SP1-NC_11_/IP6 assemblies, which exhibit similar Q155 side-chain chemical shifts. Q155 forms an N–H···O hydrogen bond with E159, and the observed shift indicates that HRF-4467 alters the local environment of this interaction. Consistent with retention of the hydrogen bond, the corresponding structural models exhibit an average Q155 Nε2–E159 O distance of approximately 3 Å across the six chains (PDB IDs: 9D6E, 7R7P, and 5I4T). Two distinct Q155 resonance patterns further reveal heterogeneity in its local environment, which is particularly pronounced in the HRF-4467-bound SP1-V7A assemblies (**Figure S6d**). In addition, ^1^H-^13^C HETCOR spectra of the HRF-4467-bound SP1-V7A assemblies show inter-residue correlations involving the GVGGP motif and residues G156, N195, Q155, and Q219 (**Figure S7a and S7b**, Supporting Information). Together, the Q155 chemical-shift changes, heterogeneity, and additional inter-residue correlations demonstrate that HRF-4467 reorganizes and increases ordering around the IP6-binding site, with the most pronounced effects in the SP1-V7A assemblies.

Collectively, our findings identify IP6 as an active participant in the structural response to HRF-4467 binding. Although IP6 remains coordinated by the K158 and K227 lysine rings, its reduced dynamics and distinct conformations in WT and SP1-V7A assemblies reveal its conformational adaptability within the 6HB. Together with the observed changes in junction-helix dynamics, these findings support the notion that HRF-4467 stabilizes the CA-SP1 region in the BVM-resistant SP1-V7A variant. This coupling among inhibitor binding, IP6 conformation, and junction-helix dynamics provides a possible structural basis for HRF-4467’s retained antiviral activity against SP1-V7A.

### Allosteric effects of HRF-4467 in WT and BVM-resistant variant

The structural perturbations induced by HRF-4467 extend beyond the immediate inhibitor-binding pocket, affecting neighboring intermolecular interfaces, while preserving the overall architecture of the immature Gag lattice. To further characterize these effects, we compared CP and rINEPT-(H)NH spectra of HRF-4467-bound WT and SP1-V7A assemblies (**Figure 4f**). The rINEPT-(H)NH spectra selectively detect dynamic regions of the protein^57^. Prominent amide resonances assigned to K158 near the IP6-binding site, SP1-T10, and other residues in the CA_CTD_ and SP1 regions were observed in the HRF-4467-bound WT assemblies but were substantially attenuated or absent in the SP1-V7A assemblies, indicating reduced dynamics in the variant. These observations are consistent with increased ordering at the CA-SP1 junction and around the IP6-binding site in the SP1-V7A assemblies. Dynamic differences were also evident at the hexamer–hexamer interface (**Figure 4f, 4g**), where residues W184 and M185 exhibited more restricted motion in the HRF-4467-bound SP1-V7A assemblies than in HRF-4467-bound WT assemblies. These findings suggest that the effects of the V7A polymorphism in the inhibitor-bound assemblies extend beyond the site of the polymorphism to interhexamer contacts within the immature lattice.

## DISCUSSION

Our findings show that HRF-4467 stabilizes the CA-SP1 region in both WT and BVM-resistant SP1-V7A assemblies, consistent with its ability to potently inhibit HIV-1 isolates containing the SP1-V7A polymorph^35^. This behavior aligns with the broader antiviral activity against Gag polymorphic variants reported for structurally related second-generation BVM-derived MIs, including GSK232, GSK-8999, and related compounds a and b, which incorporate structural modifications that extend the BVM scaffold (**Figure 1 and Table 1**)^22, 58, 59^. GSK-8999 exhibited activity against 57 of 62 HIV-1 isolates^22^, while GSK232 retained activity against multiple Gag polymorphic variants and subsequently advanced to clinical evaluation^58, 59^. These observations suggest that modifying the BVM scaffold can reduce the impact of CA-SP1 sequence variation on inhibitor activity. Our MAS NMR structures reveal an extended inhibitor-protein interaction network that supports 6HB stabilization in SP1-V7A, providing a structural basis for HRF-4467’s retained antiviral activity against this polymorphic variant^35^. These findings emphasize that extending and modifying the BVM scaffold may preserve inhibitor-mediated 6HB stabilization across Gag variants. In this context, comparison of HRF-4467 with the structurally related inhibitor GSK232 highlights differences in charge, conformational flexibility, and lipophilicity that may influence their interactions with the CA-SP1 binding pocket. As described above, GSK232 contains a C-28 alkylamine substituent with two sp³-hybridized tertiary amines. These groups are expected to undergo protonation under physiological conditions, giving GSK232 greater cationic character and a higher propensity to adopt zwitterionic states than HRF-4467. Although their common BVM-derived scaffold may favor broadly similar binding orientations, the distinct geometry and conformational flexibility of the GSK232 C-28 substituent could alter the orientation of its chlorobenzyl group.

The predicted cLogP values indicate that the *para-*chloro analog of HRF-4467 is more lipophilic (cLogP=9.3) than GSK232 (cLogP=8.5), whereas HRF-4467(F) has a predicted lipophilicity (cLogP=8.75) comparable to that of GSK-232. Thus, replacing chlorine with fluorine modestly reduces the predicted lipophilicity of the HRF-4467 scaffold. Further reductions may be achieved by incorporating a GSK3640254-inspired C-3 substituent (cLogP=7.9) or an additional carboxylate-containing (1*S*)-1-(fluoromethyl)cyclohex-4-ene-1,2-dicarboxylate moiety (cLogP=5.2). These modifications offer testable strategies to tune the physicochemical properties of HRF-4467, although experimental evaluation is needed to assess their effects on antiviral potency and pharmacokinetic behavior. Beyond the inhibitor’s physicochemical properties, sequence variation within the CA-SP1 binding site may influence susceptibility to BVM-derived MIs. Notably, HIV-2 and related SIVs exhibit intrinsic resistance to GSK232^34^, potentially reflecting sequence differences within the CA-SP1 binding site. Specifically, the HIV-1 CA-SP1 junction (CA-V230 to SP1-T8) comprises the sequence VLAEAMSQVT, whereas the corresponding region in HIV-2 (CA-L230 to SP1-I8) is LMAEALKEVI (PDB: 8FZC). These substitutions substantially alter the local amino acid composition of the MI-binding region and could modify its geometry, hydrophobicity, and electrostatic environment, thereby weakening interactions with BVM-derived MIs. However, the structural and molecular basis underlying this resistance remains to be established.

As summarized in **Table 1**, compounds a, b, and GSK8999 exhibit comparable antiviral activity against WT HIV-1, whereas compound a shows markedly reduced activity against the SP1-V7A variant. This difference suggests that the dimethyl substitution adjacent to the chlorophenyl group (**Figure 1**) contributes to activity against SP1-V7A. Stereochemistry at this position also strongly influences activity: the analog bearing the (*R*)-methyl substituent is substantially more potent against QVT polymorphs than the corresponding (*S*)-methyl diastereomer^22^. The resistance of SP1-V7A to BVM may, in part, result from altered hydrophobic packing within the 6HB, as substituting valine with alanine removes the two Cγ methyl groups of the V7 side chain and leaves only the Cβ methyl group of alanine. MIs bearing extended substituents may compensate for the resulting loss of hydrophobic packing within the 6HB. In HRF-4467, the geminal dimethyl groups of the isobutyramide are linked to the chlorophenyl group through a chlorobenzamide-containing extension and may form additional hydrophobic contacts within the pocket. They may also restrict rotation relative to the corresponding des-methyl analogs. These interactions could reduce dependence on the SP1-V7 side chain and thereby stabilize the V7A 6HB. The carbonyl group of the cyclopentenone ring, oriented toward SP1-T8 and SP1-A11, may provide additional stabilizing interactions in both WT and V7A assemblies. These structure-activity relationships suggest that coordinated optimization of hydrophobic packing, stereochemistry, and the C-28 and C-3 substituents may facilitate the development of MIs with improved activity against CA-SP1 polymorphic variants.

## CONCLUSIONS

Our MAS NMR study provides an atomistic structural basis for HRF-4467 activity by defining its binding mode within CA_CTD_-SP1-NC_11_/IP6 assemblies. HRF-4467 occupies an extended binding pocket that spans the canonical CA-SP1 6HB and extends toward the SP1-NC interface, where its interactions stabilize and order the junction helices. ^19^F MAS NMR spectroscopy, combined with protein-embedded QM calculations, reveals conformational heterogeneity of the bound inhibitor, indicating that HRF-4467 adopts multiple conformations within the 6HB. Beyond the immediate binding pocket, HRF-4467 induces structural and dynamic changes throughout the assembly while preserving the overall architecture of the immature lattice. Inhibitor binding restricts the microsecond-timescale dynamics of IP6, which adopts distinct conformations in WT and SP1-V7A assemblies, revealing structural coupling among HRF-4467, the CA-SP1 6HB, and IP6. The extended C-28 modification of HRF-4467 may provide additional stabilizing interactions that reduce dependence on the SP1-V7 side chain, allowing 6HB stabilization to be retained in the BVM-resistant SP1-V7A variant. Together, these findings provide a molecular basis for HRF-4467’s retained activity against SP1-V7A and establish a structural framework for developing next-generation HIV-1 MIs with broader activity against CA-SP1 polymorphic variants.

## Supplementary Information

The Supplementary information contain additional figures showing electrostatic potential maps and conformational analysis of HRF-4467 and its fluoro analog, a ^19^F CPMAS NMR spectrum of CA_CTD_-SP1-NC_11_/HRF-4467(F)/IP6 assemblies, 2D CORD spectra of CA_CTD_-SP1-NC_11_/IP6, CA_CTD_-SP1-NC_11_/HRF-4467/IP6, CA_CTD_-SP1/BVM/IP6, and CA_CTD_-SP1(V7A)-NC_11_/HRF-4467/IP6 assemblies, rINEPT-TOBSY spectra of CA_CTD_-SP1-NC_11_/HRF-4467/IP6, CA_CTD_-SP1/BVM/IP6, and CA_CTD_-SP1(V7A)-NC_11_/HRF-4467/IP6 assemblies, CP-(H)NH spectra of CA_CTD_-SP1-NC_11_/IP6, CA_CTD_-SP1-NC_11_/HRF-4467/IP6, CA_CTD_-SP1/BVM/IP6, and CA_CTD_-SP1(V7A)-NC_11_/HRF-4467/IP6 assemblies, HC HETCOR spectra of CA_CTD_-SP1-NC_11_/IP6, CA_CTD_-SP1-NC_11_/HRF-4467/IP6, and CA_CTD_-SP1(V7A)-NC_11_/HRF-4467/IP6 assemblies, and structural ensembles of HRF-4467-bound WT and SP1-V7A CA_CTD_-SP1-NC_11_ assemblies in the presence of IP6, and ^19^F solution NMR spectra of free HRF-4467 (F). The supplementary tables provide summaries of MAS NMR restraints, experimental and DFT-calculated ^19^F chemical shifts, HRF-4467 (F) conformational analysis, and NMR structure statistics.

## ACKNOWLEDGMENTS

This work was supported by the National Institutes of Health (NIH Grant 1U54AI170791 to A.M.G., T.P., B. G.-P.). We acknowledge the support of the National Science Foundation (NSF Grant CHE0959496) for the acquisition of the 850 MHz NMR spectrometer, and NIH 1S10OD025185-01A1 for funding the upgrade of the 600 MHz solution NMR spectrometer at the University of Delaware. Work in the Freed lab is supported by the Intramural Research Program of the Center for Cancer Research, National Cancer Institute, National Institutes of Health and by funding from the NIH Office of AIDS Research. We thank Kumar Tekwani Movellan and Changmiao Guo for helpful discussions on NMR experiments.

## Author contributions

T.P., A.M.G., B.K.G.-P., E.O.F., and O.P. conceived the project. T.N.P., J.A.M., K.K., and R.Z. prepared the samples. K.K., T.P., C.M.Q., and R.Z. conducted the investigation, performed NMR experiments, and curated the data. K.K. performed the computations and formal analysis. T.P., A.M.G., B.K.G.-P., C.M.Q., E.O.F., S.D.A., T.J.N., and C.T.W. provided resources. T.P., A.M.G., B.K.G.-P., and O.P. supervised the project. K.K. and T.P. wrote the original draft. All authors reviewed and edited the manuscript and approved the final version.

## COMPETING INTERESTS

The authors declare that they have no competing interests.

## DATA AVAILABILITY

Data supporting the findings are provided in the paper and/or the Supplementary Materials. Atomic coordinates have been deposited in the Protein Data Bank (PDB) under accession codes **39MG** and **39MI**. The MAS NMR chemical shifts have been deposited in the Biological Magnetic Resonance Bank (BMRB) under entry IDs **31317** and **31318**. Other supporting data and scripts for structure calculations, analysis of calculation results, and structure visualization are available from the corresponding authors upon request. Materials used in this study are available from the corresponding authors, subject to applicable material transfer agreements or other restrictions.

## Notes

### Competing Interest Statement

The authors have declared no competing interest.

