## Supplementary Information for "Structural Basis of Second-Generation HIV-1 Maturation Inhibitor Activity by Magic Angle Spinning NMR"

[1]Department of Chemistry and Biochemistry, University of Delaware, Newark, Delaware, USA; [2]Pittsburgh Center for HIV Protein Interactions, University of Pittsburgh School of Medicine, Pittsburgh, Pennsylvania, USA; [3]Department of Biochemistry, University of Utah, Salt Lake City, Utah, USA; [4]HIV Dynamics and Replication Program, Center for Cancer Research, National Cancer Institute, Frederick, Maryland, USA; [5]DFH Pharma, Gaithersburg, Maryland, USA; [6]Department of Structural Biology, University of Pittsburgh School of Medicine, Pittsburgh, Pennsylvania, USA.

**Current address:**
† T.P.: Department of Chemistry, University of California, Berkeley, Berkeley, CA, 94720, USA.

***Corresponding authors:** Tatyana Polenova, Department of Chemistry, University of California, Berkeley, CA, USA,; Angela M. Gronenborn, Department of Structural Biology, University of Pittsburgh School of Medicine, 3501 Fifth Ave., Pittsburgh, PA 15260, USA,; Barbie K. Ganser-Pornillos, Department of Biochemistry, University of Utah, Salt Lake City, Utah, USA,.

### Table of Contents


- **Figure S1.** Conformational landscape of HRF-4467(F).
- **Table S1.** Summary of the predicted $^{19}F$ chemical shifts for HRF-4467(F) conformations.
- **Figure S2.** Experimental $^{19}F$ chemical shifts of free HRF-4467 (F)
- **Table S2.** Experimental $^{19}F$ chemical shifts and predicted isotropic magnetic shielding constants.
- **Table S3.** Summary of Structure Statistics for WT $CA_{CTD}$-SP1-$NC_{11}$/HRF-4467/IP6 and $CA_{CTD}$ SP1(V7A)-$NC_{11}$/HRF-4467/IP6 assemblies.
- **Table S4.** Summary of MAS NMR Restraints for $CA_{CTD}$-SP1-$NC_{11}$/HRF-4467/IP6 assemblies.
- **Figure S3.** MAS NMR structural ensemble of the $CA_{CTD}$-SP1-$NC_{11}$/HRF-4467/IP6 and $CA_{CTD}$-SP1(V7A)-$NC_{11}$/HRF-4467/IP6 crystalline assemblies showing the central hexamer.
- **Figure S4.** DFT-calculated electrostatic potentials of HRF-4467 and HRF-4467(F).
- **Figure S5.** Selected region of 2D CORD spectra of $CA_{CTD}$-SP1-$NC_{11}$/IP6, $CA_{CTD}$-SP1-$NC_{11}$/HRF-4467/IP6, $CA_{CTD}$-SP1/BVM/IP6 and $CA_{CTD}$-SP1(V7A)-$NC_{11}$/HRF-4467/IP6 assemblies.
- **Figure S6.** Selected region of 2D rINEPT-TOBSY, $^{1}H$-detected 2D CP-(H)NH and 2D NCACX MAS spectra showing HRF-4467-induced changes in SP1-tail dynamics and CA-Q155 side-chain conformation.
- **Figure S7.** Selected region of $^{13}C$-detected 2D HC HETCOR spectra showing HRF-4467-induced changes in the local environment of CA-Q155 and CA-E159.

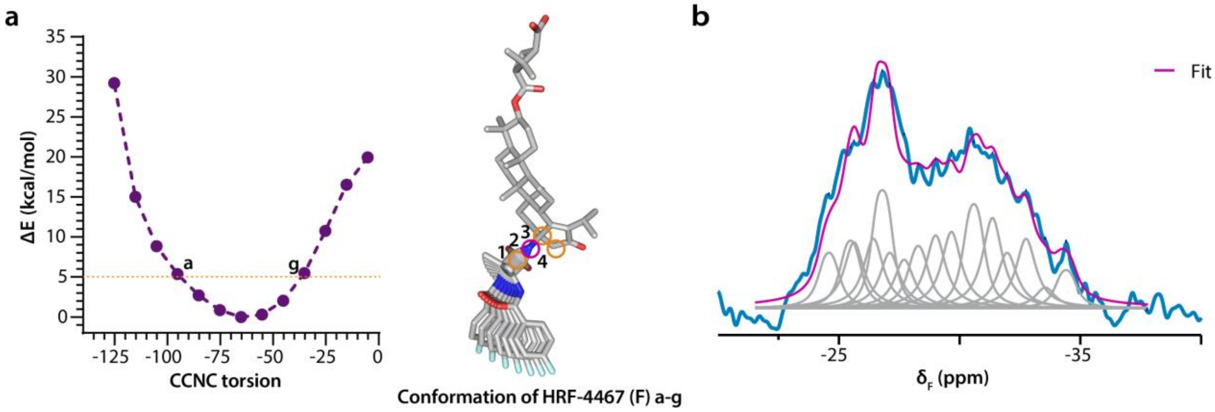


**Figure S1 | Conformational landscape of HRF-4467(F). (a)** Relative conformational energy as a function of the CNCC dihedral angle (defined by atoms 1–2–3–4). Conformations within 5 kcal/mol of the global minimum are indicated by a dotted line, and their corresponding molecular structures are shown. **(b)** $^{19}$F CPMAS NMR spectrum of $CA_{CTD}$-SP1-$NC_{11}$/HRF-4467(F)/IP6 assemblies. Deconvolution of the $^{19}$F spectrum highlights the relative populations of distinct bound conformational states of HRF-4467(F). The number of components was estimated from resolved and partially resolved features in the 1D $^{19}$F CPMAS spectrum and the 2D $^{1}$H–$^{19}$F HETCOR spectrum.

**Table S1 | Summary of the predicted $^{19}$F chemical shifts for HRF-4467(F) conformations (a–g).** Conformations within 5 kcal/mol of the global minimum were selected from the torsional energy landscape. $^{19}$F isotropic shielding constants were calculated using DFT-GIAO methods, with and without a protein QM region. Electrostatic embedding was used for the rest of the 6HB and IP6 when QM regions were used**.** A uniform empirical offset of -6 ppm was applied to all calibrated $^{19}$F chemical shifts predicted at the B3LYP/cc-pVDZ level, based on comparison with experimental values of free HRF-4467 (F) (**Figure S2**), $\delta_{iso}$ = -33.35 ppm in 30% MeOH/20 % $D_2O$/50% $H_2O$, $\delta_{iso}$ = -33.80 ppm in 50% MeOH/20 % $D_2O$/30% $H_2O$ and $\delta_{iso}$ = -34.20 ppm in DMSO-$d_6$. The uncorrected predictions differ from the experimental values by ~6 ppm in HRF-4467 (F) bound states, within the ± 7.7 ppm range reported for ~95% of calibrated B3LYP/cc-pVDZ CF predictions[37].

| **Conformations of HRF-4467 (F)** | **CNCC torsion angle*** | **$^{19}$F isotropic shielding constant (ppm)** Free HRF-4467 (F) | **Predicted $^{19}$F chemical shift (ppm)** | **$^{19}$F isotropic shielding constant (ppm)** Protein + HRF-4467 (F) | **Predicted $^{19}$F chemical shift (ppm)** |
|---|---|---|---|---|---|
| Conformation a | -95.2 | 294.6275 | -37.4796 | 280.7211 | -24.9637 |
| Conformation b | -85.2 | 294.5678 | -37.4258 | 281.4906 | -25.6562 |
| Conformation c | -75.2 | 294.5633 | -37.4218 | 280.0403 | -24.3510 |
| Conformation d | -65.2 | 294.5670 | -37.4251 | 280.2124 | -24.5059 |
| Conformation e | -55.2 | 294.5754 | -37.4327 | 282.7550 | -26.7942 |
| Conformation f | -45.2 | 294.6143 | -37.4677 | 280.6523 | -24.9018 |
| Conformation g | -35.2 | 294.5607 | -37.4194 | 284.2908† | -28.1764 |

*CNCC torsion angle at 1, 2, 3, 4 atoms of HRF-4467 (F)
†The closest side chain was repositioned to eliminate steric overlap

Note: Application of solution-derived offset to protein-bound predictions assumes that the systematic computational bias is approximately transferable between the free and bound states.

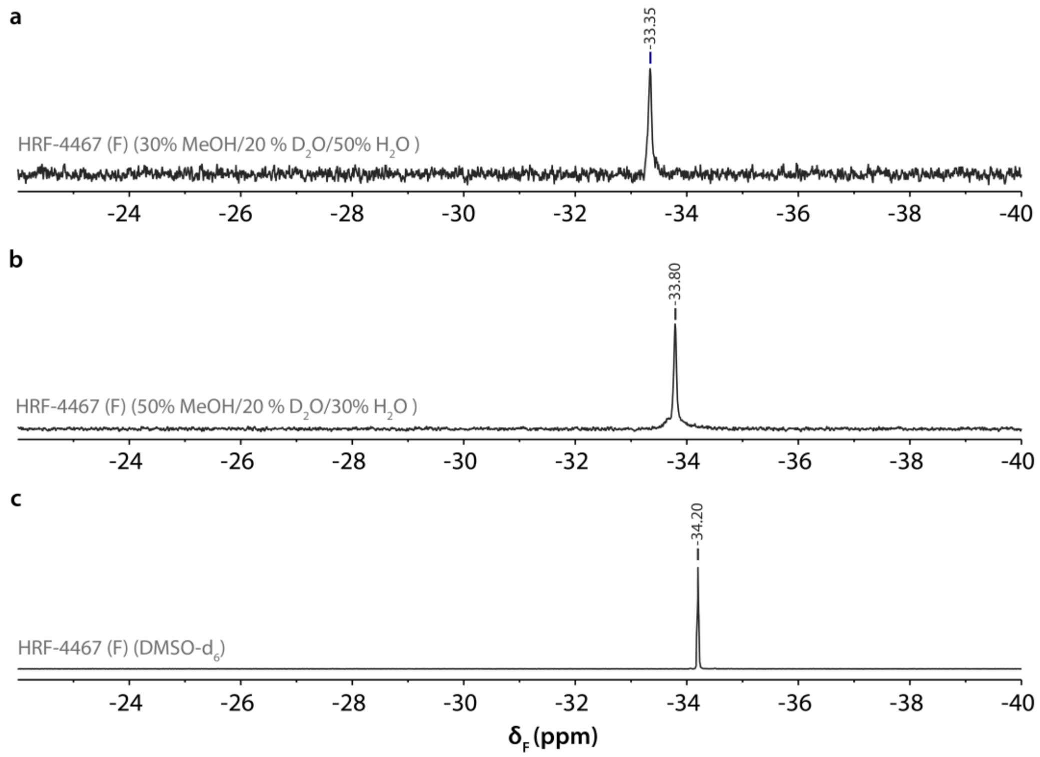


**Figure S2 | [19]F solution NMR spectra and chemical shifts of free HRF-4467 (F).** $^{19}$F NMR spectra recorded in **a)** 30% MeOH/20 % $D_2O$/50% $H_2O$, **b)** 50% MeOH/20% $D_2O$/30% $H_2O$, and **c)** DMSO-$d_6$. The corresponding chemical shifts ($\delta_{iso}$) are shown.

**Table S2 | Experimental $^{19}$F chemical shifts and predicted isotropic magnetic shielding constants.** Predictions are based on computed isotropic magnetic shielding constants and experimental chemical shifts of 707 aromatic and aliphatic CF groups[37]: $\sigma = -1.1111*\delta + 174.6514$, where δ and σ denote the chemical shift and the isotropic magnetic shielding constant, respectively.

| **Experimental $^{19}$F chemical shifts referenced to mefloquine (ppm)** | **Experimental $^{19}$F chemical shifts referenced to $CFCl_3$ (ppm)** | **Predicted $^{19}$F isotropic shielding constant (ppm) B3LYP/cc-pVDZ (ppm)** | **Corrected $^{19}$F isotropic shielding constant (ppm)** |
|---|---|---|---|
| -34.4090 | -110.9090 | 297.8824 | 291.8824 |
| -30.5026 | -107.0026 | 293.5420 | 287.5420 |
| -28.6100 | -105.1100 | 291.4391 | 285.4391 |
| -26.8490 | -103.3490 | 289.4825 | 283.4825 |
| -25.3000 | -101.8000 | 287.7614 | 281.7614 |
| -24.3700 | -100.8700 | 286.7281 | 280.7281 |

**Table S3 | Summary of structure statistics for WT $CA_{CTD}$-SP1-$NC_{11}$/HRF-4467/IP6 and $CA_{CTD}$-SP1(V7A)-$NC_{11}$/HRF-4467/IP6 assemblies.**

| | **WT $CA_{CTD}$-SP1-$NC_{11}$/HRF-4467/IP6 assemblies** | **$CA_{CTD}$-SP1(V7A)-$NC_{11}$/HRF-4467/IP6 assemblies** |
|---|---|---|
| Violations (mean ± s.d.) | | |
| Distance restraints ≥ 7.2 Å (Å) | 0.189 ± 0.006 | 0.186 ± 0.005 |
| Dihedral angle restraints ≥ 5° (°) | 2.548 ± 0.050 | 2.630 ± 0.054 |
| Max. protein-protein distance restraint violation (Å) | 2.090 | 2.040 |
| Max. protein-ligand distance restraint violation (Å) | 1.430 | 0.910 |
| Deviations from idealized geometry | | |
| Bond lengths (Å) | 0.014 ± 0.000 | 0.014 ± 0.000 |
| Bond angles (°) | 0.926 ± 0.011 | 0.912 ± 0.011 |
| Impropers (°) | 0.898 ± 0.021 | 0.898 ± 0.021 |
| Average pairwise r.m.s.d.* (Å) | | |
| Heavy | 1.34 ± 0.23 | 1.43 ± 0.17 |
| Backbone (N, Cα, C) | 0.97 ± 0.14 | 1.00 ± 0.12 |

*pairwise r.m.s. deviation was calculated among 10 lowest energy central hexamers

**Table S4 | Summary of MAS NMR restraints for $CA_{CTD}$-SP1-$NC_{11}$/HRF-4467/IP6 assemblies.**

| | | | | | |
|---|---|---|---|---|---|
| **HRF-4467-Protein restraints** | | | | | |
| $^{1}$H(HRF-4467)-$^{13}$C(protein) | | 8 | | | |
| $^{19}$F(HRF-4467)-$^{1}$H(protein) | | 8 | | | |
| **$IP_6$-Protein restraints** | | | | | |
| $^{31}$P($IP_6$)- $^{1}$H(protein) | | 26 | | | |
| **Protein restraints** | | | | | |
| **$CA_{CTD}$-SP1-$NC_{11}$/HRF-4467/$IP_6$ distance restraints** | | $^{13}$C-$^{13}$C | $^{15}$N-$^{13}$C | $^{1}$H-$^{15}$N | $^{1}$H-$^{13}$C |
| Unambiguous | Intra-residue | 724 | 283 | 81 | 144 |
| | Sequential (\|i-j\| = 1) | 491 | 117 | 7 | 6 |
| | Medium range (1<\|i-j\|<4) | 681 | 107 | 1 | 9 |
| | Long range (\|i-j\|≥5) | 139 | 16 | 0 | 2 |
| | Long range (\|i-j\|>4) (inter-chain) | 26 | 0 | 0 | 0 |
| | Long range (\|i-j\|>4) (inter-hexameric) | 14 | 0 | 0 | 0 |
| Ambiguous | Intra-residue | 22 | 0 | 6 | 0 |
| | Sequential (\|i-j\| = 1) | 94 | 4 | 0 | 0 |
| | Medium range (1<\|i-j\|<4) | 135 | 2 | 0 | 0 |
| | Long range (\|i-j\|≥5) | 20 | 0 | 0 | 0 |
| Total number of unambiguous restraints of each type | | 2075 | 523 | 89 | 161 |
| Total number of restraints | | 3131 | | | |
| Restraints/Residue | | 28 | | | |
| Percent Completeness | | 38 % and 63 % at 7.2 and 6.0 Å distance cut-off respectively (C−C only) | | | |

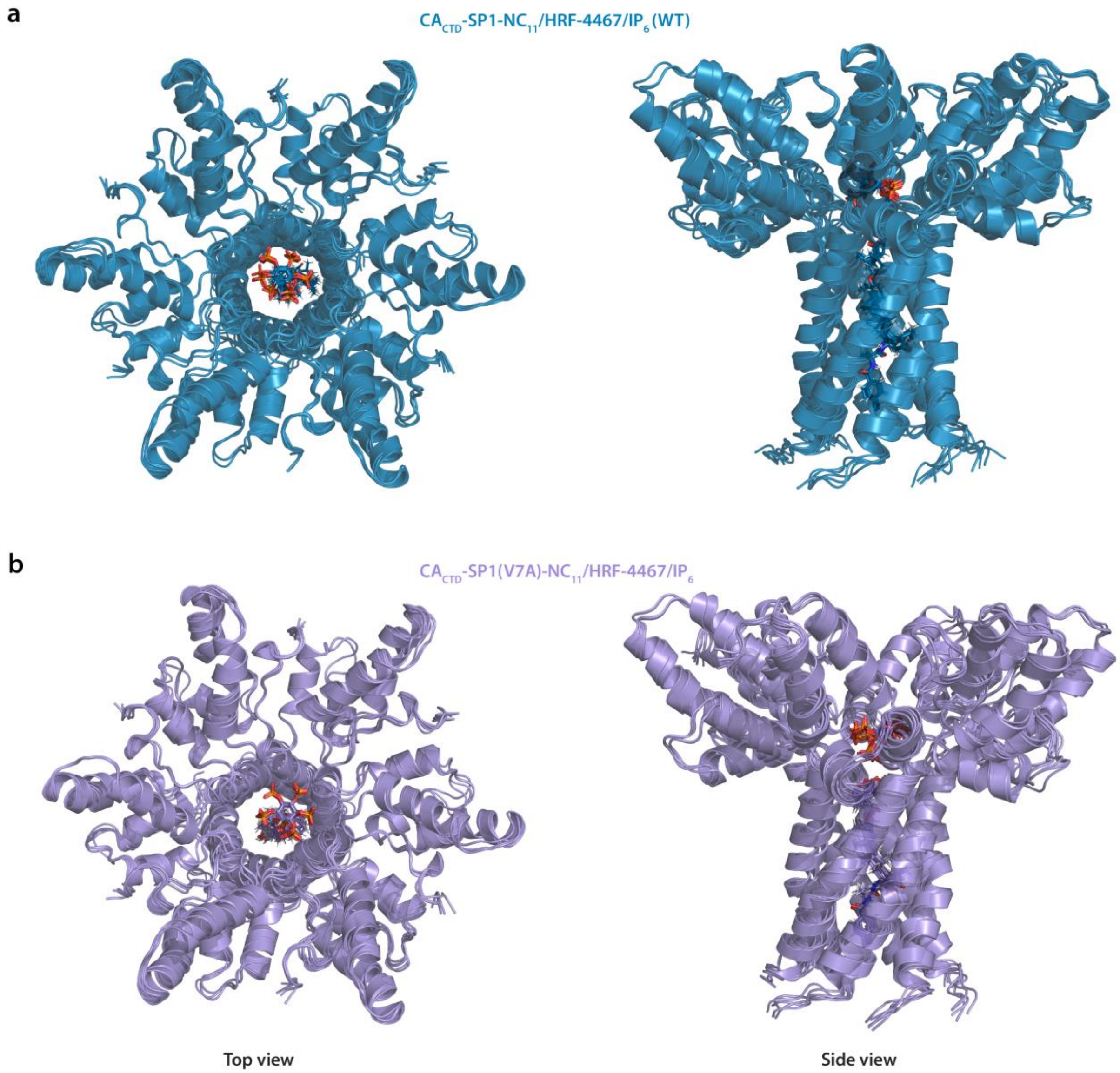


**Figure S3 | MAS NMR structural ensembles of the $CA_{CTD}$-SP1-$NC_{11}$/HRF-4467/IP6 and $CA_{CTD}$-SP1(V7A)-$NC_{11}$/HRF-4467/IP6 crystalline assemblies showing the central hexamer.**

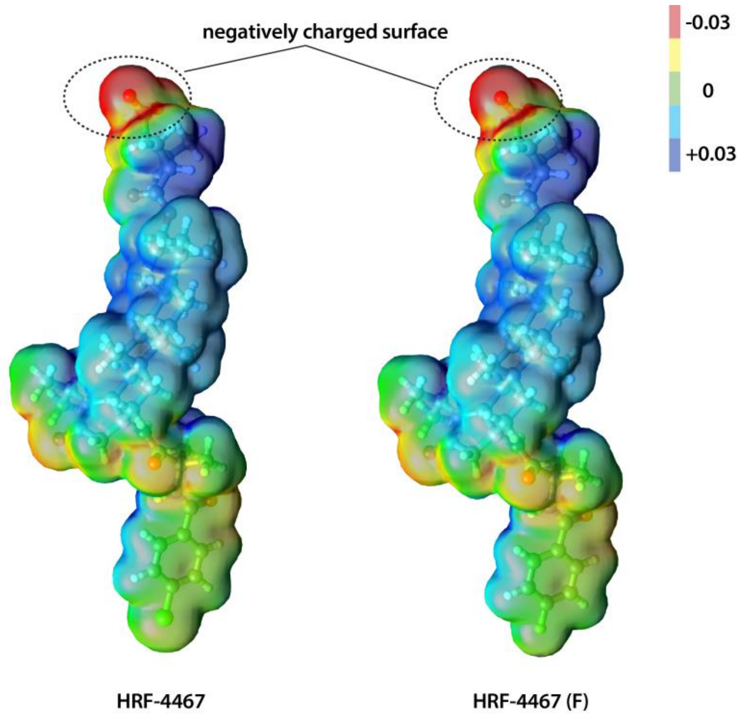


**Figure S4 | Electrostatic potentials of HRF-4467 and HRF-4467(F), calculated at the B3LYP/6-31G(d,p) level and mapped onto their electron-density isosurfaces.**

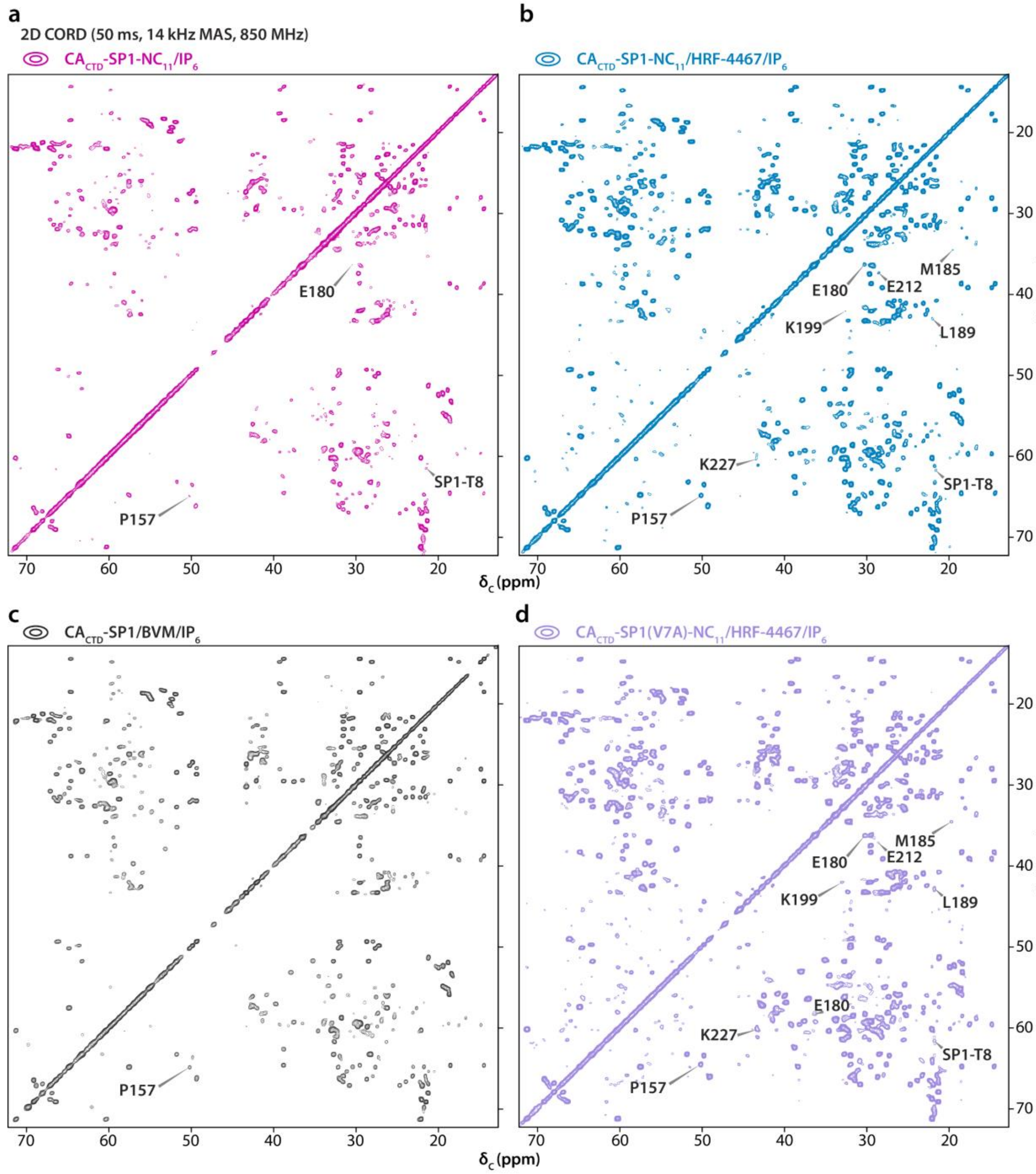


**Figure S5 | Selected regions of 2D CORD spectra** of **a)** CA$_{CTD}$-SP1-NC$_{11}$/IP6 (magenta), **b)** CA$_{CTD}$-SP1-NC$_{11}$/HRF-4467/IP6 (blue), **c)** CA$_{CTD}$-SP1/BVM/IP6 (black) and, **d)** CA$_{CTD}$-SP1(V7A)-NC$_{11}$/HRF-4467/IP6 (violet) assemblies. Spectral comparison highlights HRF-4467-induced changes in residues from helices 9, 10, and 11 at the hexamer–hexamer interface, residues near IP6 (CA-P157 and CA-K227), and the SP1-T8 residue within the 6HB.

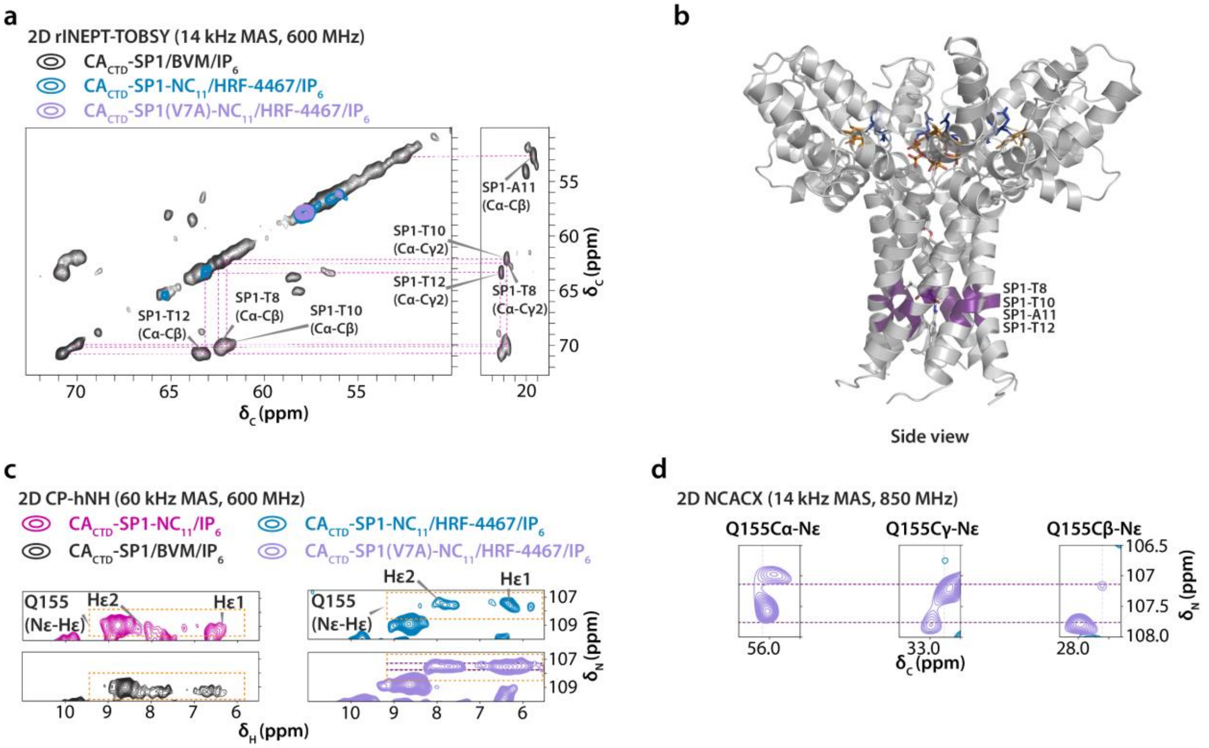


**Figure S6 | a), b)** Overlay of selected regions of 2D rINEPT-TOBSY spectra of CA$_{CTD}$-SP1/BVM/IP6 (black), CA$_{CTD}$-SP1-NC$_{11}$/HRF-4467/IP6 (blue), and CA$_{CTD}$-SP1(V7A)-NC$_{11}$/HRF-4467/IP6 (violet) assemblies, showing assignments of SP1 T8-T12 tail and corresponding structural representation is shown (the corresponding cross peaks are absent in WT CA$_{CTD}$-SP1-NC$_{11}$ and CA$_{CTD}$-SP1(V7A)-NC$_{11}$ assemblies; the SP1-T12 Cα diagonal peak is present in WT and all other diagonal peaks from SP1 tail are absent in V7A construct). The TOBSY mixing time was 7.6 ms. **c)** Selected region of $^{1}$H-detected CP-(H)NH MAS NMR correlation spectra of CA$_{CTD}$-SP1-NC$_{11}$/IP6 (magenta), CA$_{CTD}$-SP1-NC$_{11}$/HRF-4467/IP6 (blue), CA$_{CTD}$-SP1/BVM/IP6 (black), and CA$_{CTD}$-SP1(V7A)-NC$_{11}$/HRF-4467/IP6 (violet) assemblies showing the effect of HRF-4467 binding on CA-Q155 side chains (highlighted in dashed box, Hε1 and Hε2 are CA-Q155 side chain NH$_2$ protons and $^{15}$N chemical shift difference $\Delta\delta \approx 1.8$-2 ppm). **d)** Selected region of 2D NCACX spectrum and chemical shift assignment of CA-Q155 side chain.

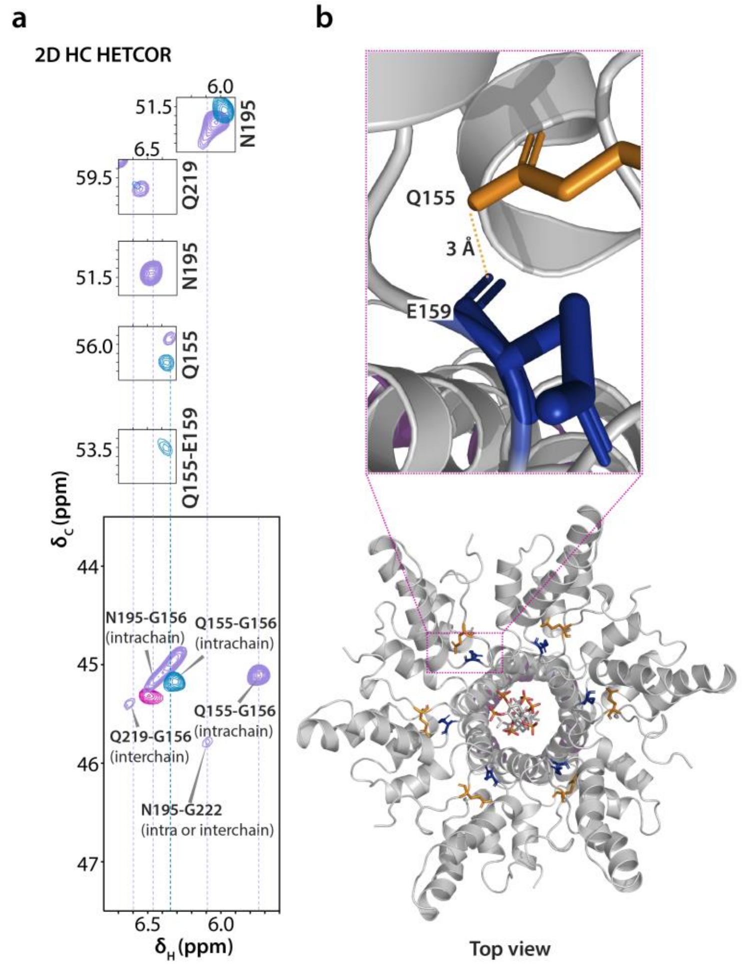


**Figure S7 | a), b)** Overlay of selected regions of $^{13}$C-detected 2D HC HETCOR correlation MAS NMR spectra of $CA_{CTD}$-SP1-$NC_{11}$/IP6 (magenta), $CA_{CTD}$-SP1-$NC_{11}$/HRF-4467/IP6 (blue), and $CA_{CTD}$-SP1(V7A)-$NC_{11}$/HRF-4467/IP6 (violet) assemblies, and structural model of $CA_{CTD}$-SP1-$NC_{11}$/HRF-4467/IP6 hexamer, and an expansion showing residues CA-Q155 (orange) and CA-E159 (dark blue).
